# Local Shifts in Inflammatory and Resolving Lipid Mediators in Response to Tendon Overuse

**DOI:** 10.1101/2021.01.08.425901

**Authors:** James F. Markworth, Kristoffer B. Sugg, Dylan C. Sarver, Krishna Rao Maddipati, Susan V. Brooks

**Affiliations:** Department of Molecular & Integrative Physiology, University of Michigan, Ann Arbor, Michigan; Department of Orthopaedic Surgery, University of Michigan, Ann Arbor, Michigan; Department of Surgery, University of Michigan, Ann Arbor, Michigan; Department of Cellular & Molecular Physiology, Johns Hopkins University, Baltimore, Maryland; Department of Pathology, Lipidomics Core Facility, Wayne State University, Detroit, Michigan; Department of Biomedical Engineering, University of Michigan, Ann Arbor, Michigan

## Abstract

Tendon inflammation has been implicated in both adaptive connective tissue remodeling and overuse-induced tendinopathy. Lipid mediators control the initiation and resolution of inflammation, but their roles within tendon are largely unknown. Here we profiled local shifts in intratendinous lipid mediators via liquid chromatography-tandem mass spectrometry in response to synergist ablation-induced plantaris tendon overuse. Sixty-four individual lipid mediators were detected in homogenates of habitually loaded plantaris tendons from healthy ambulatory rats. This included many bioactive metabolites of the cyclooxygenase (COX), lipoxygenase (LOX), and epoxygenase (CYP) pathways. Synergist ablation induced a robust inflammatory response at day 3 post-surgery characterized by epitenon infiltration of polymorphonuclear leukocytes (PMNs) and macrophages (MΦ), heightened expression of inflammation-related genes, and increased intratendinous concentrations of the pro-inflammatory eicosanoids thromboxane B_2_ (TXB_2_) and prostaglandin E_2_ (PGE_2_). By day 7, MΦ became the predominant myeloid cell type in tendon and there were further delayed increases in other COX metabolites including PGD_2_, PGF_2α_ and PGI_2_. Specialized pro-resolving mediators (SPMs) including protectin D1 (PD1) and resolvin D6 (RvD6), as well as related pathway markers of D-resolvins (17-HDoHE), E-resolvins (18-HEPE) and lipoxins (15-HETE) were also increased locally in response to tendon overuse, as were many anti-inflammatory fatty acid epoxides of the CYP pathway (e.g. EpETrEs). Nevertheless, intratendinous prostaglandins remained markedly increased even following 28 days of tendon overuse together with a lingering MΦ presence. These data reveal a delayed and prolonged local inflammatory response to tendon overuse characterized by an overwhelming predominance of pro-inflammatory eicosanoids and a relative lack of pro-resolving lipid mediators.

## Introduction

Tendons are dense bands of connective tissue responsible for transfer of force from skeletal muscle to bone (1). Like skeletal muscle, tendons can undergo compensatory hypertrophy in response to heightened mechanical loading (2). On the other hand, repetitive tendon overuse is a major contributor to the development of tendinopathy, a common degenerative condition characterized by chronic pain and loss of function (3). Inflammation occurs following either acute tendon injury (4–9) or repetitive overuse (10–17), but its role in the etiology of tendinopathy has been a matter of debate (18). The term tendinitis was classically used to describe symptoms of painful non-ruptured tendons, inferring key involvement of an inflammatory component (19). However, an apparent lack of polymorphonuclear leukocytes (PMNs) within diseased tendons led to the view that tendinopathy is rather a degenerative condition of tendinosis that is devoid of inflammation (20). Nevertheless, recent studies employing modern antibody based immunohistochemical staining techniques demonstrate that leukocytes, most notably macrophages (MΦ), are indeed often present within diseased human tendons (21), stimulating a resurgence of interest into the potential role of inflammation in tendon biology (22).

Lipid mediators are bioactive metabolites of dietary essential polyunsaturated fatty acids (PUFA), such as omega-6 (n-6) arachidonic acid (AA, 20:4n-6), as well as omega-3 (n-3) eicosapentaenoic acid (EPA, 20:5n-3) and docosahexaenoic acid (DHA, 22:6n-3) (23). A wide-range of lipid mediators can be endogenously produced via the cyclooxygenase (COX), lipoxygenase (LOX), and epoxygenase (CYP) pathways (24). These eicosanoids and docosanoids act as important autocrine/paracrine signaling molecules in a range of physiological processes, most notably in mediating the inflammatory response (25). Prior studies of tendon have focused overwhelmingly on the prostaglandins, classical eicosanoid metabolites generated via the COX-1 and −2 pathways (26). Local concentrations of prostaglandin E_2_ (PGE_2_) are well known to increase in rodent models of either acute tendon injury (27, 28) or heightened mechanical loading (28, 29), as well as within peritendinous tissues of exercising humans (30, 31). While PMNs (32) and MΦ (33) are major classical cellular sources of PGE_2_, resident tendon fibroblasts (tenocytes) can also produce and release PGE_2_ in response to either inflammatory cytokines (34) or mechanical stimulation (35).

Resolution of the acute inflammatory response, characterized by cessation of PMN influx and clearance of infiltrating leukocytes from the site of inflammation, was originally thought to be a passive event (36). More recently, distinct families of specialized pro-resolving mediators (SPMs) were shown to be produced during the resolution phase (37). SPM families identified to date include the AA-derived lipoxins (e.g. LXA_4_) (38), EPA-derived E-series resolvins (e.g. RvE1) (39), and DHA-derived D-series resolvins (e.g. RvD1) (40), protectins (e.g. PD1) (41), and maresins (e.g. MaR1) (42). Collectively these autocoids act as endogenous stop signals to limit further PMN influx (43), while simultaneously stimulating key MΦ functions required for timely resolution and tissue repair (44). The discovery of SPMs has inspired the development of novel therapeutic strategies to modulate inflammation by mechanisms that are distinct from classical anti-inflammatory approaches such as non-steroidal anti-inflammatory drugs (NSAIDs) (45). Administration of resolution agonists, termed immunoresolvents, can limit inflammation and expedite its resolution, while simultaneously relieving pain and stimulating tissue repair (46). Unlike NSAIDs that interfere with musculoskeletal tissue remodeling (26), SPMs were recently found to rather exert pro-regenerative actions following skeletal muscle injury (47, 48).

Interest into the potential role of SPMs in tendon is emerging (49). The AA-derived SPM lipoxin A4 (LXA_4_) (50), and its cell surface receptor ALX/FPR2 (51), were both found to be increased in inflamed equine tendons and isolated human tenocytes were recently shown to produce a range of different lipid mediators *in-vitro*, including both pro-inflammatory eicosanoids and SPMs (52). Exogenous SPM treatment can also suppress the release of pro-inflammatory cytokines by isolated human tenocytes *in-vitro*, indicative of a potential important role of these novel bioactive lipid mediators in controlling tendon inflammation (53–55). Therefore, the goal of the present study was to assess for the first time whether changes in mechanical loading of tendon modulates these endogenous inflammation-resolving pathways *in-vivo*.

## Methods

### Animals

Male Sprague-Dawley rats were obtained from Charles River Laboratories and housed under specific pathogen-free conditions with ad-libitum access to food and water. Rats were used for experiments at approximately 6-months of age. All animal experiments were approved by the University of Michigan Institutional Animal Care and Use committee (IACUC) (PRO00006079).

### Plantaris Tendon Overuse

Myotenectomy induced synergist ablation was used to assess the local inflammatory response to mechanical overload of the plantaris musculotendinous unit as originally described by Goldberg et al. 1967 (56). Rats were anesthetized with 2% isoflurane and preemptive analgesia provided by subcutaneous injection of buprenorphine (0.03 mg/kg) and carprofen (5 mg/kg). The skin overlying the posterior hind-limb was shaved and scrubbed with chlorhexidine and ethyl alcohol. A midline incision was made through the overlying skin and paratenon to visualize the gastrocnemius/soleus (Achilles) tendon. A full thickness tenectomy was performed to surgically remove the entire Achilles tendon mid-substance, while leaving the plantaris tendon intact. The paratenon was loosely re-approximated and the incision was closed using 4-0 Vicryl sutures. The procedure was then repeated on the contralateral limb to induce bilateral mechanical overload of both the left and right plantaris tendons. Rats were returned to their cage to recover and monitored until ambulatory with free access to food and water. Postoperative analgesia was provided via subcutaneous injection of buprenorphine (0.03 mg/kg) at 12 h post-surgery and animals were monitored daily for any signs of pain or distress for 7 days. Age matched male rats served as non-surgical ambulatory control animals for collection of habitually loaded plantaris tendons.

### Tissue Collection

Animals were euthanized via induction of bilateral pneumothorax while under isoflurane anesthesia. The plantaris musculotendinous unit was carefully dissected and a sample of the tendon mid-substance isolated by severing its distal insertion at the calcaneus and at its proximal border near the myotendinous junction. The isolated plantaris tendon samples were blotted dry, weighed, and then cut transversely with a scalpel blade into three separate pieces. The mid-portion of the plantaris tendon, allocated to immunohistochemical analysis, was oriented longitudinally on a plastic support, covered with a thin layer of optimal cutting temperature (OCT) compound, and rapidly frozen in isopentane cooled on liquid nitrogen. The remaining proximal and distal portions of the plantaris tendon, allocated to mRNA expression and LC-MS/MS analysis respectively, were weighed and then snap frozen in liquid nitrogen. Samples were stored at −80°C until further analysis. Samples from the mid-belly region of the plantaris muscles from these same rats were also collected for analysis and their mediator lipidomic profile is reported separately (48).

### Histological Analysis of Tendon Inflammation

Tissue cross-sections (10 μm) were cut at −20°C from the mid-portion of OCT embedded plantaris tendons in a cryostat (CryoStar NX50, Thermo Fisher Scientific). Sections were adhered to SuperFrost Plus slides, air dried at room temperature, fixed in ice-cold acetone at −20°C for 10 min. Following air drying to evaporate residual acetone the fixed slides were blocked for 1 h at room temperature in 10% normal goat serum (Invitrogen, Thermo Fisher Scientific, 10000C) in phosphate buffered saline (PBS) prior to overnight incubation at 4°C with blocking buffer containing primary antibodies raised against HIS48 (Abcam, Ab33760, 1:20), CD68 (Abcam, ab31630, 1:50), and CD163 (Santa Cruz, sc-33560, 1:50), to simultaneously detect rat myeloid cell populations including PMNs, ED1 (M1-like) MΦ, and ED2 (M2-like) MΦ respectively. The following day, slides were washed in PBS and then incubated for 1 h at room temperature with secondary antibodies including Goat Anti-Mouse IgG1 Alexa Fluor 488 conjugate (Invitrogen, Thermo Fisher Scientific, A21121, 1:500 in PBS), Goat Anti-Mouse IgM Alexa Fluor 555 conjugate (Invitrogen, Thermo Fisher Scientific, A21426, 1:500 in PBS), and Goat Anti-Rabbit IgG (H+L) Alexa Fluor 647 (Invitrogen, Thermo Fisher Scientific, A21245, 1:500 in PBS). Wheat germ agglutinin (WGA) CF405S conjugate (Biotium, 29027, 100 μg/mL in PBS) was also included in the secondary antibody solution in order to label and visualize the extracellular matrix. Following further washing in PBS, slides were mounted using Fluorescence Mounting Medium (Agilent Dako, S302380) and allowed to dry overnight protected from light at room temperature Fluorescent images were captured using a Nikon A1 inverted confocal microscope.

### LC-MS/MS Based Metabolipidomic Profiling of Tendon

Plantaris tendon samples were mechanically homogenized in 1 mL of phosphate buffered saline (PBS) using a bead mill. The tissue homogenates were centrifuged at 3000 × *g* for 5 min and the resulting supernatant collected. Supernatants (0.85 ml) were spiked with 150 μl methanol containing 5 ng each of 15(S)-HETE-d8, 14(15)-EpETrE-d8, Resolvin D2-d5, Leukotriene B4-d4, and Prostaglandin E1-d4 as internal standards for recovery and quantitation and mixed thoroughly. The samples were then extracted for PUFA metabolites using C18 extraction columns as previously described (48, 57, 58). Briefly, the internal standard spiked samples were applied to conditioned C18 cartridges, washed with 15% methanol in water followed by hexane and then dried under vacuum. The cartridges were eluted with 2 x 0.5 ml methanol with 0.1% formic acid and then then eluate was dried under a gentle stream of nitrogen. The residue was re-dissolved in 50 μl methanol-25 mM aqueous ammonium acetate (1:1) and subjected to LC-MS analysis. HPLC was performed on a Prominence XR system (Shimadzu) using Luna C18 (3 μm, 2.1 × 150 mm) column as previously described (59, 60). The HPLC eluate was directly introduced to electrospray ionization source of a QTRAP 5500 mass analyzer (ABSCIEX) in the negative ion mode and monitored by a Multiple Reaction Monitoring (MRM) method to detect unique molecular ion – daughter ion combinations for each of the lipid mediators using a scheduled MRM around the expected retention time for each compound. Spectra of each peak detected in the scheduled MRM were recorded using Enhanced Product Ion scan to confirm the structural identity. The data were collected using Analyst 1.7 software and the MRM transition chromatograms were quantitated by MultiQuant software (both from ABSCIEX). The internal standard signals in each chromatogram were used for normalization, recovery, as well as relative quantitation of each analyte.

LC-MS/MS data was analyzed using MetaboAnalyst 4.0 (61). Analytes with >50% missing values were removed from the data set and remaining missing values were replaced with half of the minimum positive value in the original data set. Heat maps were generated in MetaboAnalyst using the Pearson distance measure and the Ward clustering algorithm following auto scaling of features without data transformation. Volcano and PCA plots were generated in R following Log2 data transformation using the EnhancedVolcano and FactoMineR packages respectively

### RNA Extraction and RT-qPCR

Tendon samples were homogenized for 45 seconds at 4 m/s in 600 μL TRIzol reagent using a Fisherbrand™ Bead Mill 4 Homogenizer (Thermo Fisher Scientific, 15-340-164) with reinforced 2 mL tubes (Thermo Fisher Scientific, 15-340-162) and 2.4 mm metal beads (4 beads/tube) (Thermo Fisher Scientific, 15-340-158). RNA was isolated by Phenol/Chloroform extraction and RNA yield determined using a NanoDrop Spectrophotometer (Nanodrop 2000c, Thermo Fisher Scientific). Genomic DNA was removed by incubation with DNase I (Ambion, Thermo Fisher Scientific, AM2222) followed by its heat inactivation. Total RNA (250 ng) was then reverse transcribed to cDNA using SuperScript™ VILO™ Master Mix (Invitrogen, 11-755-050) and RT-qPCR performed on a CFX96 Real-Time PCR Detection System (Bio-Rad, 1855195) in duplicate 20 μL reactions of iTaq™ Universal SYBR® Green Supermix (Bio-Rad, 1725124) with 1 μM forward and reverse primers (Table 1). Relative mRNA expression was determined using the 2^-ΔΔCT^ method with *B2m* serving as an endogenous control.

**Table 1:**
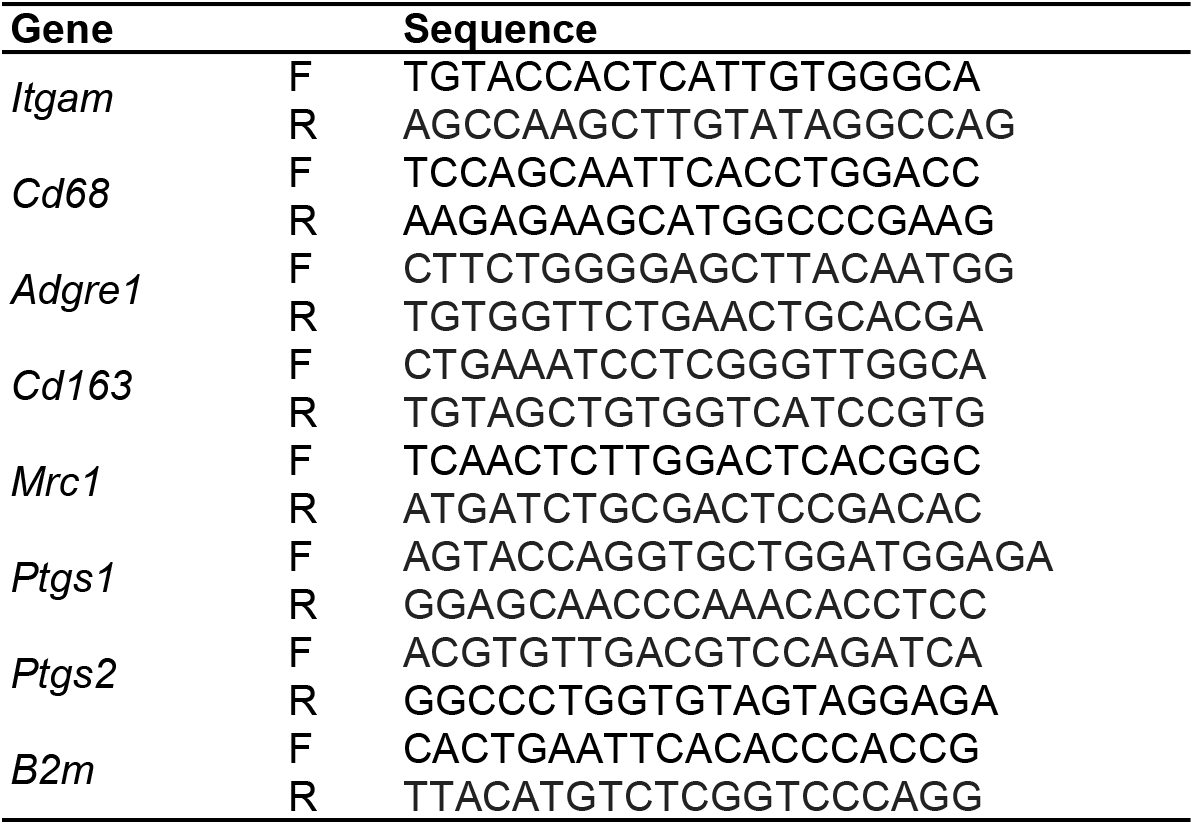
Real-time reverse transcription PCR primers

### Statistics

Data is presented as the mean ± SEM with raw data from each individual tendon sample shown. Statistical analysis was performed in GraphPad Prism 7. Between group differences were tested by two-tailed unpaired students t-tests (2 groups) or by a one-way analysis of variance (ANOVA) followed by pair-wise Holm-Sidak post-hoc tests (≥3 groups). For time-course experiments, multiple comparison testing was made compared to a single control group of tendons obtained from ambulatory control rats that did not undergo surgery. p≤0.05 was used to determine statistical significance.

## Results

### Lipid Mediator Profile of Tendon

We initially examined the basal lipid mediator profile of habitually loaded tendons via LC-MS/MS based targeted metabolipidomics. A total of sixty-four individual lipid mediator species were reliably detected (signal to noise ratio >3 and peak quality >0.2 in at least 50% of samples) in plantaris tendon homogenates from rats undergoing normal cage activity (Figure 1A). These included many bioactive metabolites of n-6 AA derived via the COX-1 and 2 pathways [prostaglandins, e.g. PGD_2_, PGE_2_, PGF_2α_, PGI_2_ (measured as its inactive non-enzymatic hydrolysis product 6-keto-PGF1_α_) and thromboxanes [e.g. TXA_2_ (measured as its inactive non-enzymatic hydrolysis product TXB_2_) and the related thromboxane synthase metabolite 12-hydroxy-heptadecatrienoic acid (12-HHTrE)] (Figure 1A, Supplemental Table 1A). Monohydroxylated AA metabolites of the three major mammalian lipoxygenase enzymes (5-, 12, and 15-LOX) were also detected including 5-, 12-, 15-hydroxy-eicosatetraenoic acids (HETEs)] (Figure 1A, Supplemental Table 1B). Finally, AA metabolites of the epoxygenase (CYP 450) pathway were present in tendon including 5(6)-, 8(9)-, 11(12)-, 14(15)-epoxy-eicosatrienoic acid regioisomers (EpETrEs) (Figure 1A, Supplemental Table 1C). In addition to these eicosanoids, several metabolites of the parent n-6 PUFA linoleic acid (LA, 18:2n-6) were highly abundant in tendon homogenates, including those derived via both the LOX pathway [9- and 13-hydroxy-octadecadienoic acid (HODEs) and downstream oxo-octadecadienoic acids (OxoODEs)] and CYP pathway [9(10)- and 12(13)-epoxy-octadecenoic acids (EpOMEs) and downstream dihydroxy-octadecenoic acids (DiHOMEs)] (Figure 1A).

**Figure 1.**
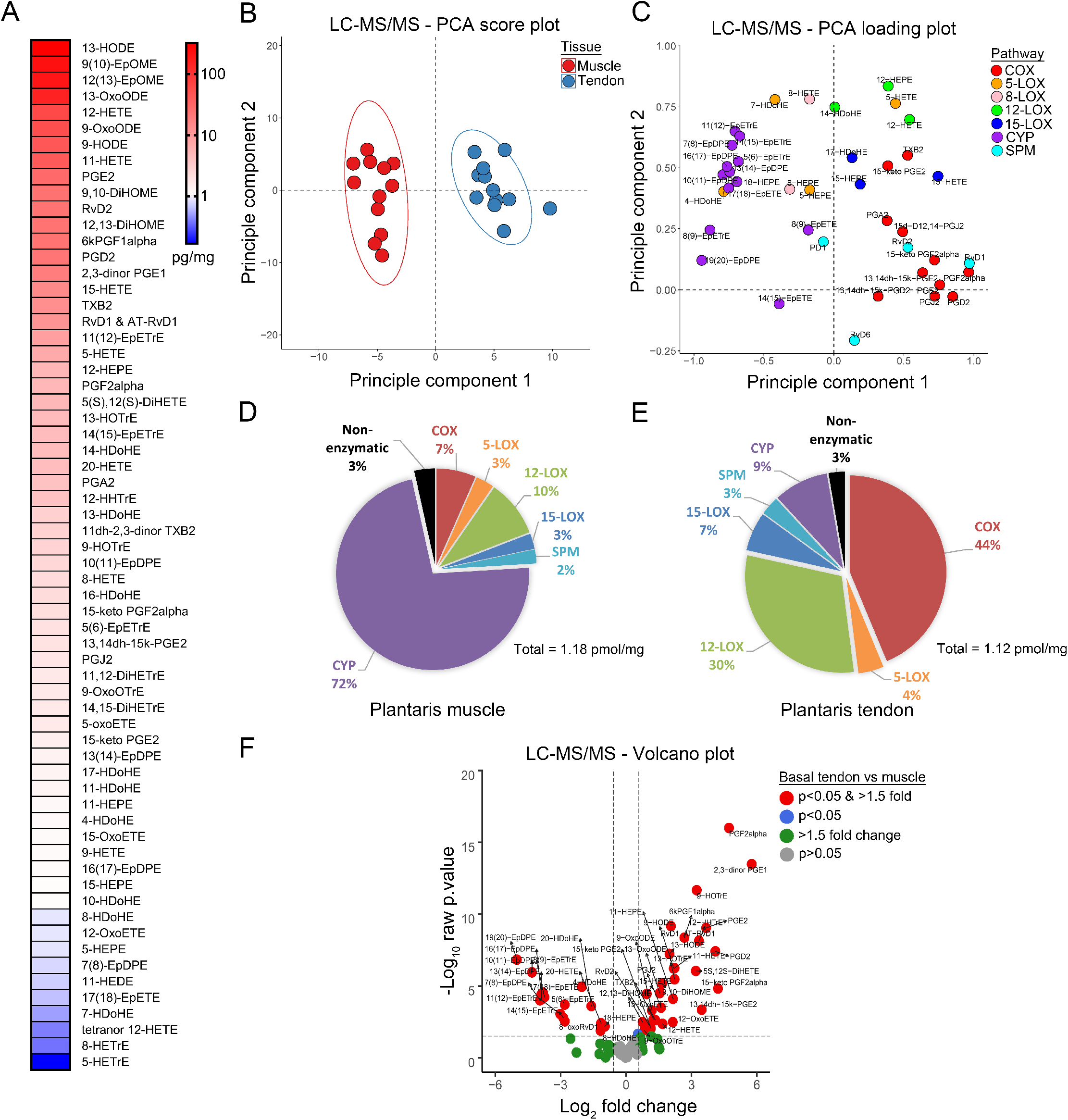
Divergent lipid mediator profiles of functionally related musculoskeletal tissues. **A:** Complete profile of lipid mediators detected by tandem liquid chromatography-mass spectrometry (LC-MS/MS) analysis of plantaris tendon homogenates from healthy rats undergoing habitual cage activity ranked by absolute concentration. **B:** Unsupervised principle component analysis (PCA) score plots of the overall LC-MS/MS profile of functionally associated plantaris tendon and muscle samples. **C:** PCA loading plot showing the relative contributions of some representative analytes from each major enzymatic biosynthetic pathway. **D:** Percentage composition of the overall mediator lipidome of plantaris muscle samples. **E:** Percentage composition of the overall mediator lipidome of plantaris tendon samples. **D-E:** Linoleic acid (18:2n-6) metabolites (e.g. HODEs & EpOMEs) are excluded from graphical presentation and are shown in Supplemental Table 1. **F:** Volcano plot showing the direction, magnitude, and statistical significance of lipid mediator concentrations between tendon and muscle. Each dot represents a single analyte, positive Log2 fold changes (FC) indicate lipid mediator concentrations which were enriched in tendon, and negative Log2 FC indicate those enriched in muscle. Analytes with +1.5 absolute FC (+0.58 Log2 FC) or −1.5 FC (−0.58 Log2 FC) between tissue type and unadjusted p<0.05 were considered to differ significantly between tendon and muscle samples. P-values were determined by two-tailed unpaired t-tests.

Many n-3 PUFA metabolites were also detected in tendon, albeit generally at relatively lower concentrations than the above n-6 PUFA products. This included the DHA-derived SPM, resolvin D1 (RvD1), as well as 17-hydroxy-docosahexaenoic acid (17-HDoHE), the primary intermediate 15-LOX metabolite of n-3 DHA produced during the initial step of D-series resolvin biosynthesis. Other SPMs including the E-series resolvins (e.g. RvE1), protectins (e.g. PD1), and maresins (e.g. MaR1) were generally below the limits of detection in resting tendon homogenates (Supplemental Table 1D). Nevertheless, key pathway markers in the biosynthesis of the E-series resolvins [(18-hydroxy-eicosapentaenoic acid (18-HEPE)] and maresins [14-hydroxy-docosahexaenoic acid (14-HDoHE)] were detected. Finally, CYP pathway derived epoxides of n-3 PUFAs including EPA [5(6)-, 8(9)-, 11(12)-, 14(15)-, 17(18)-epoxy-eicosatetraenoic acid (EpETEs)] and DHA [7(8)-, 10(11)-, 13(14)-, 16(17)-, 19(20-epoxy-docosapentaenoic acid (EpDPE)] were also present within habitually loaded tendons. These data reveal a wide range of novel bioactive lipid mediators within healthy tendon tissue *in-vivo* for the first time.

### Functionally Associated Musculoskeletal Tissues Exhibit Highly Distinct Metabolipidomic Profiles

The complete LC-MS/MS profile of plantaris skeletal muscle samples from these same rats has been previously published (48). Unsupervised principle component analysis (PCA) revealed that the mediator lipidome of the tendon samples analyzed here was highly distinct from that of the habitually loaded plantaris muscles from these same rats (Figure 1B). The corresponding loading plots displaying some representative lipid mediators from each major enzymatic biosynthetic pathway that defined the distinct lipid mediator profiles of muscle and tendon samples are shown in Figure 1C. When pooled over these major biosynthetic pathways, 72% of the overall mediator lipidome of muscle comprised CYP pathways metabolites (e.g. EpETrEs, EpETEs, and EpDPEs), with only 7%, 10%, and 3% of total metabolites derived from the COX (e.g. PGE_2_), 12-LOX (e.g. 12-HETE), and 15-LOX (e.g. 15-HETE) pathways respectively (Figure 1D). In contrast, the tendon mediator lipidome overwhelmingly comprised COX (42%), 12-LOX (28%), and 15-LOX (6%) pathway metabolites, with only 9% of total lipid mediators derived from the CYP pathway (Figure 1D). Despite these differences in the relative composition of mediator lipidome between tissues, total lipid mediator concentration was similar between muscle and tendon when normalized to tissue mass (~1.2 pmol/mg) (Figure 1 D-E).

Parametric statistical analysis revealed that of the total ninety-four individual lipid mediators that were detected in either tendon or muscle tissue, thirty-five were significantly enriched in tendon (p<0.05 and >1.5-fold), while seventeen were significantly enriched in muscle (p<0.05 and >1.5-fold) (Figure 1F). When compared to muscle, tendon contained relatively lower absolute concentrations of anti-inflammatory CYP pathway metabolites including many epoxide products of AA [5(6)-, 8(9)-, 11(12)-, and 14(15)-EpETrEs], EPA [8(9)-, 14(15)- & 17(18)-EpETEs], and DHA [7(8)-, 10(11)-, 13(14)-, 16(17)-, and 19(20)-EpDPEs]. 20-HETE, a ω-hydroxylase CYP metabolite of AA was similarly lacking in tendon. Finally, the primary n-3 EPA product produced during biosynthesis of the E-series resolvins, 18-HEPE, which is endogenously derived via the CYP pathway (62), was also significantly lower in tendon than muscle. On the other hand, tendon contained far higher concentrations than muscle of many COX pathway metabolites including the major AA-derived thromboxanes (TXB_2_ and 12-HHTrE) and prostaglandins (PGD_2_, PGE_2_, PGF_2α_, 6-keto-PGF1_α_). Many downstream secondary and tertiary prostaglandin metabolites of the 15-hydroxy-prostaglandin dehydrogenase (15-PGDH) and 15-oxo-prostaglandin Δ^13^-reductase pathways including 15-keto PGE_2_, 15-keto PGF_2α_, 13,14-dihydro-15-keto PGE_2_, 13,14-dihydro-15-keto PGD_2_, and 11-dihydro-2,3-dinor TXB_2_ were also enriched in tendon, as were the cyclopentenone prostaglandins including PGJ_2_, Δ^12^-PGJ_2_, and 15-deoxy-Δ^12,14^-PGJ_2_ (15d-PGJ_2_) (which are non-enzymatically derived from PGD_2_). Many LOX-pathway metabolites including 5-, 11-, 12-, and 15-HETEs, 12- and 15-oxoETEs, 11- and 12-HEPEs, 9- and 13-HODEs, 9- and 13-HOTrEs, and 5S,12S-DiHETE were also relatively enriched in tendon. Consistently, the DHA-derived SPM RvD1, which is produced by the sequential action of the 15- and 5-LOX pathways, was detected in habitually loaded tendons, but was below the limits of detection in muscle (as previously reported) (48).

### Local Shifts in Lipid Mediator Biosynthesis in Response to Synergist Ablation-Induced Tendon Overuse

In order to examine local shifts in lipid mediator biosynthesis in response to tendon overuse, we surgically removed the gastrocnemius/soleus (Achilles) tendon to induce compensatory mechanical overload upon the synergistic plantaris musculotendinous unit. Functionally overloaded plantaris tendons were then collected for analysis by LC-MS/MS at 3-, 7-, and 28-days following synergist ablation surgery (Figure 2).

**Figure 2.**
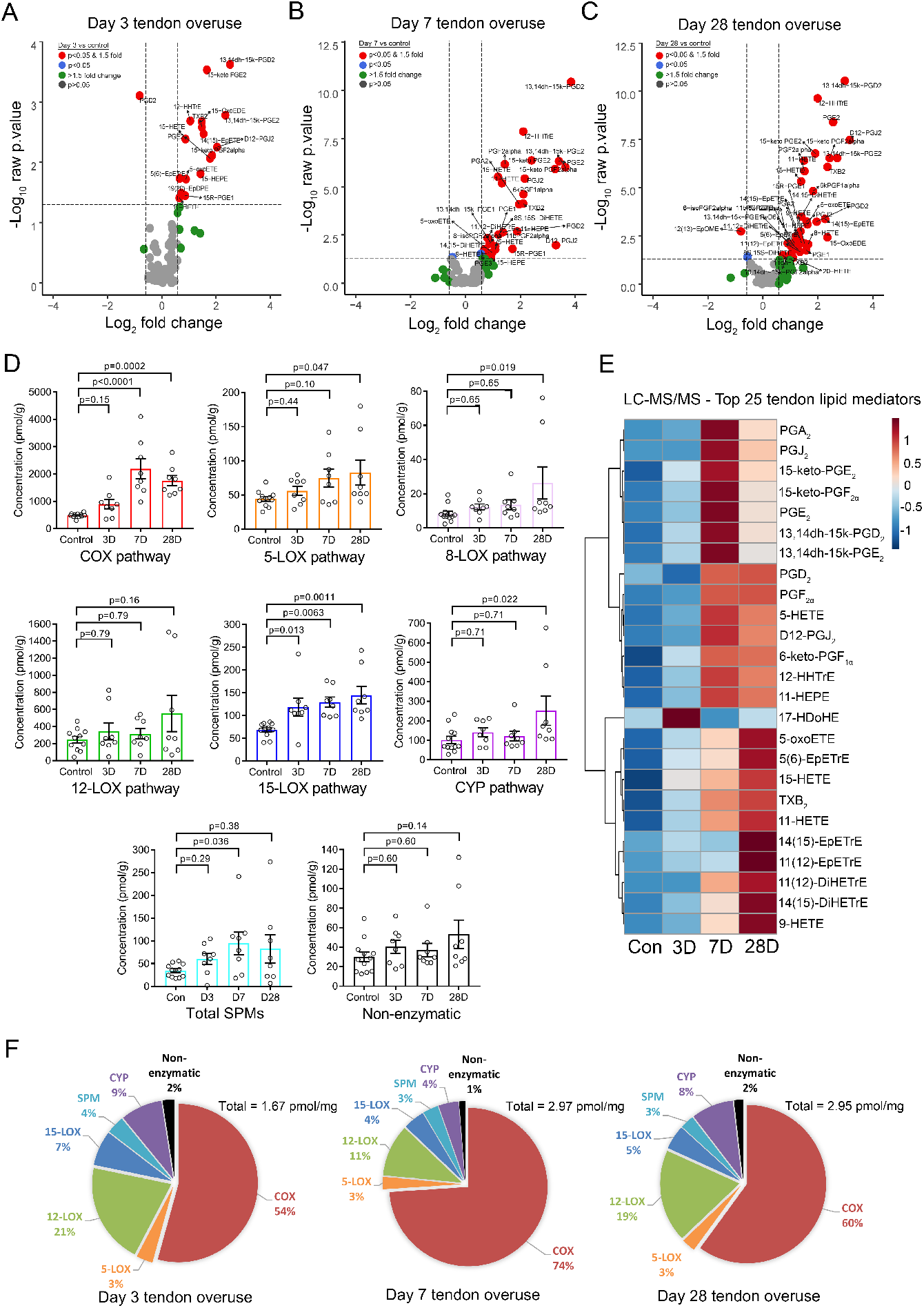
Dynamic changes in lipid mediators in response to tendon overuse. Changes in intratendinous concentrations of each individually detected lipid mediator species in overloaded plantaris tendons when compared to plantaris tendons obtained from ambulatory control rats at (**A**) day 3, (**B**) day 7, and (**C**) day 28 of recovery from synergist ablation-induced plantaris tendon overuse. **D:** Kinetics of changes in pooled lipid mediator concentration in the overloaded plantaris tendon over the time-course of recovery from synergist ablation surgery. **E:** Kinetics of changes in intratendinous concentrations of the top 25 individual lipid mediator species modulated by tendon overuse. The data for the full panel of analytes monitored by the LC-MS/MS assay is shown in Supplemental Table 1. **F:** Changes in the relative contribution of major biosynthetic pathways to the overall tendon mediator lipidome following synergist ablation induced-tendon overuse. **D-F:** Linoleic acid (18:2n-6) metabolites (e.g. HODEs & EpOMEs) are excluded from graphical presentation and are shown in Supplemental Table 1. P-values were determined by two-tailed unpaired t-tests (panel A-C) or one-way ANOVA followed by Holm-Šidák post hoc tests (panel D).

When compared to control plantaris tendons obtained from age and sex matched ambulatory rats, intratendinous concentrations of eighteen individual lipid mediator species were significantly modulated at day 3 of synergist ablation-induced plantaris tendon overuse (Figure 2A). This included increased concentrations of the COX/thromboxane synthase products TXB_2_ and 12-HHTrE, as well as the COX/prostaglandin E synthase product, PGE_2_, as well as its downstream enzymatic inactivation products 15-keto PGE_2_ and 13,14-dihydro-15-keto PGE_2_. The COX/prostaglandin D synthase product PGD_2_ was simultaneously reduced by 50%, while COX/prostaglandin F and I synthase products PGF_2α_ and PGI_2_ (measured as 6-keto-PGFiα) remained unchanged in parallel at this time-point. Primary 15-LOX metabolites of AA (15-HETE) and EPA (15-HEPE) were additionally increased at day 3 post surgery, with a similar non-significant trend also seen for 17-HDoHE (p=0.057), the analogous 15-LOX product of n-3 DHA. In contrast, most major metabolites of the 5-LOX, 12-LOX, and CYP pathways remained unchanged in tendon at day 3 following synergist ablation.

A total of thirty lipid mediators were significantly modulated following 7 days of tendon overuse (Figure 2B). This included further increases in many of the same lipid mediators seen at day 3 (e.g. TXB_2_, PGE_2_, 15-HETE) as well as additional more delayed increases in PGD_2_ and its downstream enzymatic inactivation product 13,14-dihydro-15-keto PGD_2_. Similarly, PGF_2α_ and its downstream enzymatic inactivation products 15-keto-PGF_2α_ and 13,14-dihydro-15-keto PGF_2α_ were increased, as was the fifth primary prostanoid PGI_2_ (measured as 6-keto-PGF1_α_). Finally, the Series-J cyclopentenone prostaglandins, including PGJ_2_, D12-PGJ_2_, and 15d-PGJ_2_, were produced together with their precursor PGD_2_ at day 7 following synergist ablation.

By 28-days of tendon overuse, a total of thirty-eight individual lipid mediators differed significantly from ambulatory control tendons (Figure 2C). This included persistent elevation of all major COX-metabolites (TXB_2_, PGE_2_, PGD_2_, PGF_2α_, and 6-keto-PGF_1α_) and their respective downstream metabolic inactivation products of the 15-PGDH pathway (e.g. 15-keto and 13,14-dihydro-15-keto PGs). The primary 5-LOX metabolite of n-6 AA, 5-HETE, and its metabolite 5-oxoETE, also exhibited a delayed increase at this time-point, although related 5-LOX metabolites derived from the downstream LTA4 hydrolysis pathway, including LTB4 and 12-oxoLTB4, remained below the limits of detection. Finally, CYP pathway derived epoxides of n-6 AA including 5(6)-, 11(12)-, and 14(15)-EpETrE were increased at day 28 of tendon overuse, as were some analogous CYP metabolites of EPA [11(12)-, 13(14)-, and 17(18)-EpETEs].

The temporal shifts in absolute tendon lipid mediator concentrations when pooled over major enzymatic biosynthetic pathways are summarized in Figure 2D and the time-course kinetics of the top 25 individual lipid mediator species that were modulated by tendon overuse are shown in Figure 2E. The entire quantitative tendon LC-MS/MS data set for each individual lipid mediator profiled is shown in Supplemental Table 1. Significant increases over time were found for pooled metabolites of the COX, 8-LOX, 15-LOX, and CYP pathway, as well as for pooled concentrations of detected bioactive SPMs (Figure 2D). A similar trend was also observed for pooled metabolites of the 5-LOX pathway, although this failed to reach statistical significance (p=0.076, Figure 2D). Despite a more delayed increase in local concentrations of lipid mediators with anti-inflammatory and pro-resolving actions, the overwhelming predominance of biosynthesis of classical pro-inflammatory COX pathway metabolites in mechanically overloaded plantaris tendon resulted in a progressive reduction in the percentage of the overall mediator lipidome that was derived from the LOX, CYP and SPM pathways over time (Figure 2F). Because of this, the proportion of the overall mediator lipidome consisting of classical pro-inflammatory eicosanoids derived from the COX pathway increased from 42% in control plantaris tendons from ambulatory rats (Figure 1E) to encompass 54%, 74% and 60% of the mediator lipidome at day 3, 7 and 28 of tendon overuse respectively (Figure 2F).

### Local Leukocyte Responses to Tendon Overuse

To investigate the relationship between shifts in intratendinous lipid mediator concentrations as determined by LC-MS/MS based profiling and cellular inflammation of tendon, we performed immunohistochemical staining of cross-sections of ambulatory control and mechanically overloaded plantaris tendons with antibodies to detect infiltrating myeloid cell populations including PMNs (HIS48^+^ cells), inflammatory ED1 monocytes/MΦ (CD68^+^ cells), and resident/M2-like ED2 MΦ (CD163^+^ cells) (Figure 3A). Control plantaris tendons from ambulatory rats were apparently devoid of resident leukocytes based on cellular expression of these markers. In contrast and as previously reported (48), many resident ED2 MΦ (CD68^-^CD163^+^ cells) were seen scattered throughout the internal connective tissues (e.g. perimysium and endomysium) of plantaris muscle cross-sections from these same rats. These data suggest that unlike functionally associated skeletal muscle, tendon does not appear to possess a substantial resident myeloid cell population.

**Figure 3.**
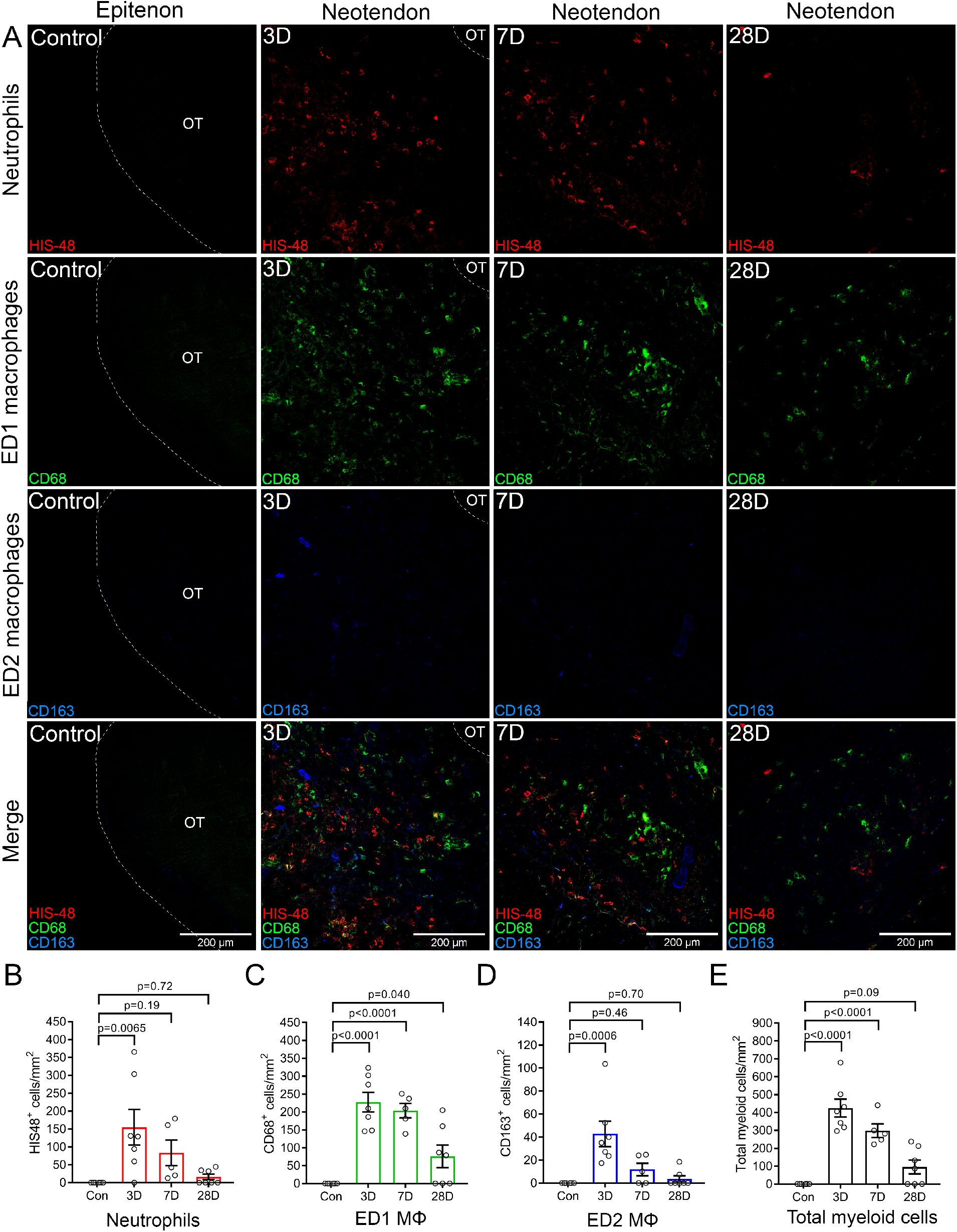
Peritendinous infiltration of inflammatory cells in response to synergist ablation-induced tendon overuse. **A:** Mechanically overloaded plantaris tendons were collected from Sprague Dawley rats at day 3, 7, and 28 following synergist ablation surgery. Plantaris tendons from ambulatory age and gender matched rats served as non-surgical controls. Tissue cross-sections were cut from the tendon mid-substance and stained with antibodies against polymorphonuclear cells (PMNs, HIS48^+^), inflammatory ED1 monocytes/macrophages (MΦ, CD68^+^), and resident/M2-like ED2 MΦ (CD163^+^). Images were captured from the periphery of control plantaris tendons (epitenon region), or from within the center of the expanded peritendinous tissue layer at the periphery of overloaded plantaris tendons (neotendon matrix). Quantification of peritendinous infiltration of (**B)** PMNs (HIS48^+^ cells), (**C)** ED1 MΦ (CD68^+^ cells), (**D)** ED2 MΦ (CD163^+^ cells), and (**E)** Total myeloid cells (sum of PMNs, ED1 MΦ and ED2 MΦ). Values are mean ± SEM of 5-7 plantaris tendon per time-point with dots representing data from each individual tendon. P-values were determined by one-way ANOVA followed by Holm-Šidák post hoc tests. The original tendon (OT) is traced with a dotted white line.

At day 3 of tendon overuse there was an accumulation of many PMNs (HIS48^+^ cells) (Figure 3A and 3B) and ED1 MΦ (CD68^+^CD163^-^ cells) (Figure 3A and 3C) throughout the expanded tissue space at the plantaris tendon periphery. ED1 MΦ were also be seen within the peripheral edges of the dense tendon core at this time-point, appearing to originate from within the epitenon (Figure 4A). A more modest increase in the histological presence of ED2 MΦ (CD68^-^CD163^+^ cells) was also seen throughout the tendon periphery, but not the tendon core, at this time-point (Figure 3A and 3D). By day 7 of tendon overuse, there was a robust histological presence of large numbers of ED1 MΦ (CD68^+^CD163^-^ cells) throughout the newly forming connective tissue layer that surrounded the original tendon (e.g. the neotendon matrix), although ED2 MΦ were no longer significantly increased (Figure 3A). Many ED1 MΦ (but not ED2 MΦ) were also seen scattered throughout the dense original tendon core at day 7 following synergist ablation (Figure 4A). By day 28 of tendon overuse, few infiltrating myeloid cells remained, in particular within the tendon core (Figure 4A). Nevertheless, some CD68^+^CD163^-^ cells could still be seen scattered throughout the peripheral neotendon matrix in the majority of samples analyzed (Figure 3C). Only a very small proportion of the CD68^+^ cells within mechanically overloaded tendons co-expressed the CD163 antigen at all time-points between day 3 and 28 of tendon overuse (Figure 3A, 3D and 4A).

**Figure 4.**
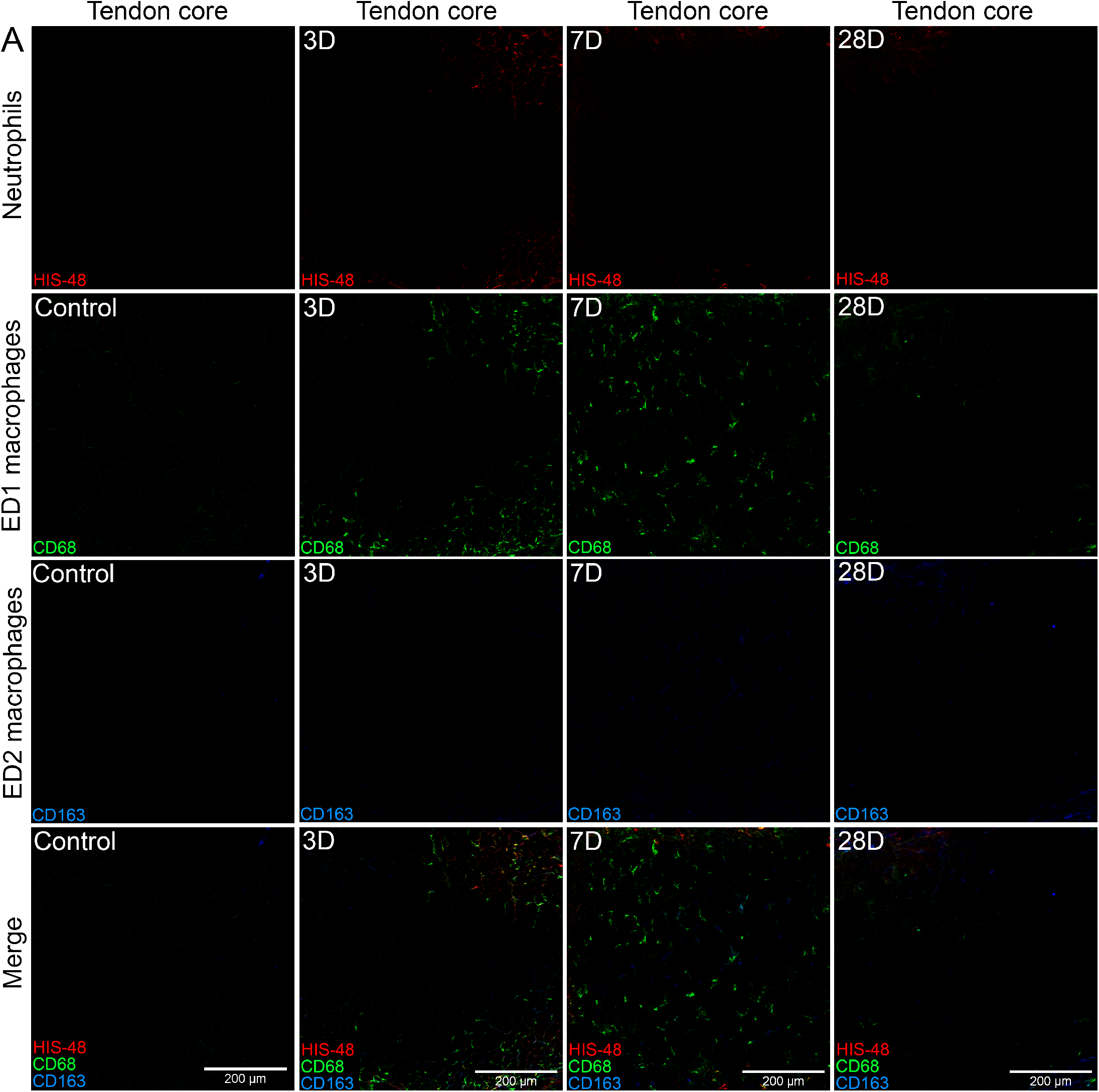
Inflammatory macrophage infiltration of the original tendon core following plantaris overuse. **A:** Mechanically overloaded plantaris tendons were collected from Sprague Dawley rats at day 3, 7, and 28 following synergist ablation-induced tendon overuse. Plantaris tendons from ambulatory age and gender matched rats served as non-surgical controls. Tissue cross-sections were cut from the tendon mid-substance and stained with antibodies against polymorphonuclear cells (PMNs, HIS48^+^ cells), inflammatory ED1 monocytes/macrophages (MΦ, CD68^+^ cells), and resident/M2-like ED2 MΦ (CD163^+^ cells). Representative images were captured from the center of the dense original tendon core.

### Expression of Inflammation-Related Genes in Tendon Following Synergist Ablation

Despite the apparent lack of histological presence of resident myeloid cells, mRNA encoding the general myeloid cell marker CD 11b *(Itgam)*, inflammatory monocyte/MΦ markers CD68 *(Cd68)* and EMR1 (the rat analog of F4/80, *Adgre1*), as well as the resident/M2-like MΦ markers CD163 *(Cd163)* and CD206 *(Mrc1)* were expressed at detectable levels in resting plantaris tendon (Figure 5). Following synergist ablation surgery, tendon mRNA expression of CD11b was increased 10-fold above ambulatory control tendons at day 3, remained elevated by 3.5 fold at day 7, but no longer differed from control levels by day 28 (Figure 5A). Similarly, expression of the MΦ markers CD68 (Figure 5B) and F4/80 (Figure 5C) both increased 15-fold and 10-fold respectively at day 3, remained increased by 3.5-fold at day 7, but had returned to basal levels by day 28 of recovery. CD163 mRNA did not differ significantly from control tendons at any time-point, but was 3.5 fold higher at day 3 of tendon overuse when compared to day 28 of recovery (p=0.023) (Figure 5D). The alternate M2-like MΦ marker CD206 (*Mrc1*) was also increased 5-fold at day 3 following synergist ablation, but no longer differed from control tendons at either day 7 or 28 (Figure 5E). Expression of both the constitutive COX-1 (*Ptgs1*) and inducible COX-2 (*Ptgs2*) isoform mRNA was detectable in ambulatory control plantaris tendons. At day 3 of tendon overuse, COX-2 mRNA expression was increased by 4-fold (Figure 5G), but COX-1 mRNA remained unchanged (Figure 5F). Neither COX-1 nor COX-2 mRNA expression differed significantly from control tendons by day 7 or 28 of continued tendon overuse.

**Figure 5.**
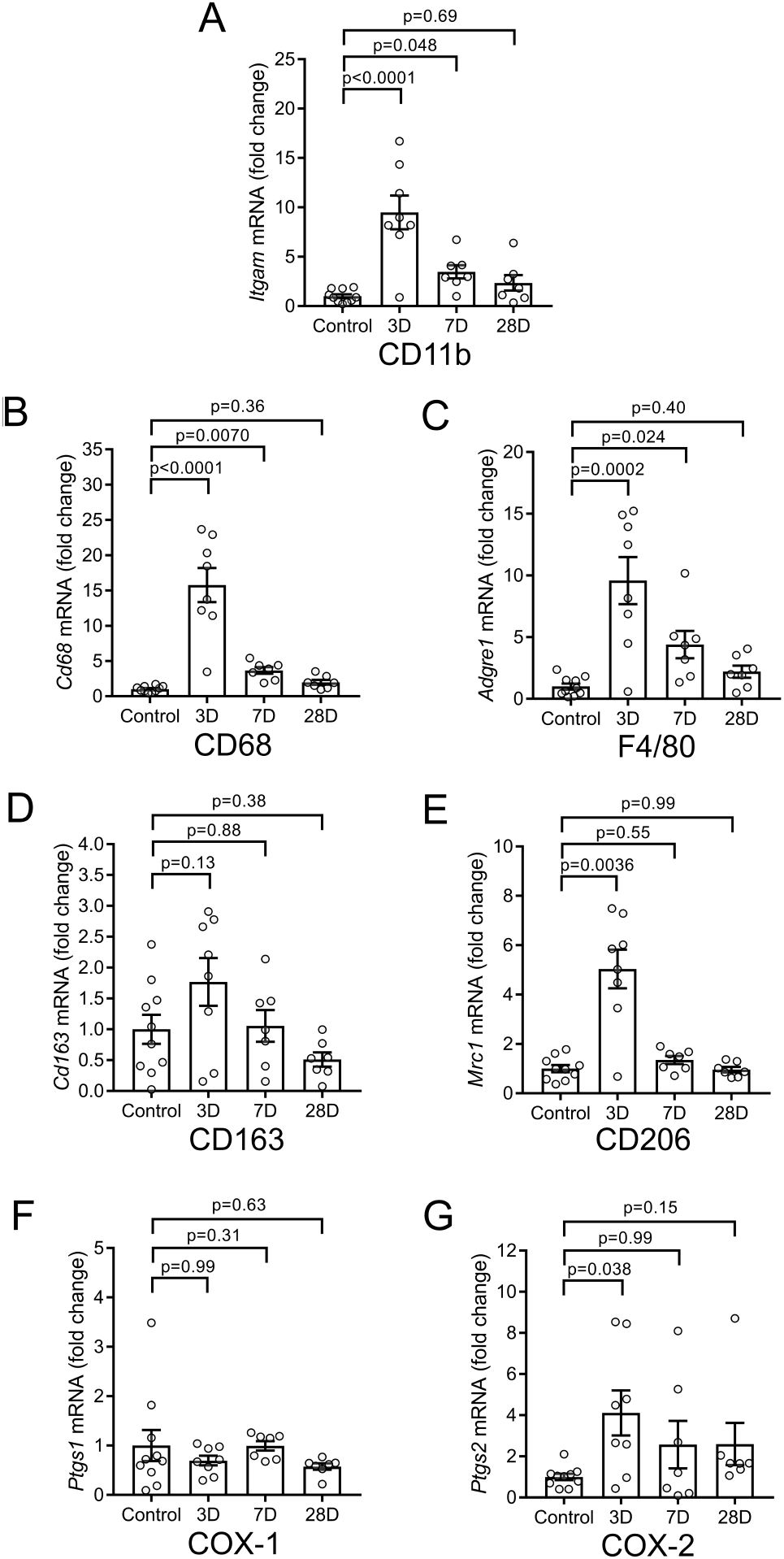
Local expression of inflammation-related genes in response to synergist ablation-induced plantaris tendon overuse. Mechanically overloaded plantaris tendons were collected from male Sprague Dawley rats at day 3, 7, and 28 following synergist ablation surgery. Habitually loaded plantaris tendons from ambulatory age and gender matched rats served as non-surgical controls. Total tendon RNA was extracted, reverse transcribed to cDNA, and expression of inflammation-related genes measured by real-time quantitative reverse transcription PCR (RT-qPCR). Relative mRNA expression (fold change from control) was determined for (**A**) CD11b (*Itgam)*, (**B**) CD68 (*Cd68)*, (**C**) EMR1 (rat analog of F4/80, *Adgre1)*, (**D**) CD163 (*Cdl63)*, (**E**) CD206 (*Mrc1)*, (**F**) cyclooxygenase-1 (COX-1, *Ptgs1)*, and (**G**) cyclooxygenase-2 (COX-2, *Ptgs2).* Beta-2-Microglobulin (*B2m*) served as an endogenous control for normalization of genes of interest. Bars show the mean ± SEM of 8-12 plantaris tendons from 4-6 rats per group with dots representing data from each individual tendon sample. P-values were determined by one-way ANOVA followed by Holm-Šidák post hoc tests.

### Changes in Tendon Mass and Total RNA Content in Response to Mechanical Overuse

Total RNA concentration (ng/mg of tissue) of the plantaris tendon was increased by 3.5 fold at 3 days of overuse, while tendon mass remained unchanged (Table 2). This resulted in a 3-fold increase in the total RNA content of the overloaded plantaris tendon (μg RNA/tendon). Both intratendinous RNA concentration and total tendon RNA content remained increased at seven days of overuse, while tendon mass was still not yet significantly altered (1.7-fold, p=0.17). By 28 days following synergist ablation a significant increase in the mass of the overloaded plantaris tendon was observed (2.2-fold). At this time-point, intratendinous RNA concentration no longer differed from that of ambulatory control tendons, but the total RNA content of the overloaded plantaris tendon remained increased by approximately 3-fold. These data show clear changes in the total RNA content of the mechanically overloaded plantaris tendon which accompany the increased cellularity resulting from leukocyte infiltration and likely also tenocyte proliferation. On this basis, it is important to consider the relative changes in mRNA expression of genes of interest (at an equivalent RNA input) in light of the marked changes in tendon total RNA content that occur following synergist ablation.

**Table 2:**
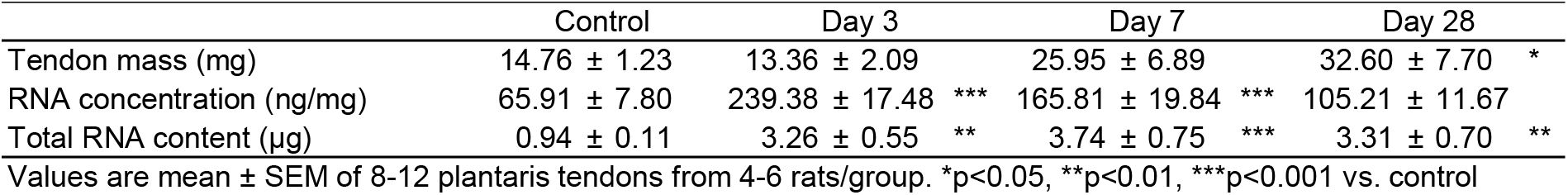
Changes in tendon mass and RNA content in response to overuse

|  | Control | Day 3 | Day 7 | Day 28 |  |
| --- | --- | --- | --- | --- | --- |
| Tendon mass (mg) | 14.76 ± 1.23 | 13.36 ± 2.09 | 25.95 ± 6.89 | 32.60 ± 7.70 | * |
| RNA concentration (ng/mg) | 65.91 ± 7.80 | 239.38 ± 17.48 *** | 165.81 ± 19.84 *** | 105.21 ± 11.67 |  |
| Total RNA content (µg) | 0.94 ± 0.11 | 3.26 ± 0.55 ** | 3.74 ± 0.75 *** | 3.31 ± 0.70 ** |  |

Values are mean ± SEM of 8-12 plantaris tendons from 4-6 rats/group. \*p<0.05, \*\*p<0.01, \*\*\*p<0.001 vs. control

## Discussion

Here we profiled local changes in lipid mediator biosynthesis following synergist ablation-induced plantaris tendon overuse. A wide range of bioactive metabolites of the COX, LOX, and CYP pathways were detected in tendon for the first time. When compared to skeletal muscle, tendons were enriched in classical pro-inflammatory eicosanoid metabolites of the COX and 12-LOX pathways, but relatively lacking in CYP-pathway derived anti-inflammatory lipid epoxides. Three days of tendon overuse induced a robust local inflammatory response characterized by heightened biosynthesis of PGE_2_ and TXB_2_, increased expression of inflammation-related genes, and peritendinous infiltration of both PMNs and MΦ. There was more delayed production of PGD_2_, PGF_2α_, 6-keto-PGF1_α_ at day 7 at which time MΦ became the predominant myeloid cell type within tendon. Proresolving lipid mediator biosynthesis was also increased following tendon overuse; however, there was a persistent reduction in the ratio of SPMs and their related LOX-derived pathway markers relative to the COX-derived prostaglandins over time. This overwhelming predominance of pro-inflammatory eicosanoids in tendon was associated with incomplete resolution of inflammation even at 28 days following synergist ablation.

The marked increase in PGE_2_ in response to plantaris tendon overuse observed in the current study supports prior studies showing that local PGE_2_ concentrations increase in response to an acute bout of exercise in both mice (28, 29) and humans (30, 31). While most prior studies in tendon have focused exclusively on PGE_2_, we show for the first time that overloaded tendons also produce substantial amounts of the other three major prostaglandins, PGD_2_, PGF_2α_, and PGI_2_ (measured as 6-keto-PGF1_α_). Biosynthesis of TXA_2_, the fifth major primary bioactive metabolite of the COX pathway, was also increased in response to synergist ablation (based on measurement of TXB_2_), which is consistent with an earlier human study in which peritendinous TXB_2_ increased following a single bout of exercise (63). Major cellular sources of specific prostanoids include blood platelets (TXA_2_) (64), PMNs (TXA_2_ and PGE_2_) (65), mast cells (PGD_2_) (66) monocytes/MΦ (PGE_2_) (67), and vascular endothelial cells (PGI_2_) (68). Thus, it is likely that the PMNs and/or MΦ that accumulated in tendon in the current study contributed substantially to the intratendinous prostaglandin response to mechanical overload. Although not assayed here, mast cells have also been found to appear locally in response to tendon overuse in prior studies and may thus contribute substantially to the PGD_2_ response (11, 16). In addition to leukocytes, fibroblasts themselves can also produce prostaglandins, most notably PGE_2_ (69), but also PGI_2_ (70) and TXA_2_ (71). Indeed, resident tendon fibroblasts (tenocytes) express both COX-1 and 2 (72), enabling them to locally produce and release PGE_2_ in response to mechanical stimulation *in-vitro* (73). Consistent with our data, recent studies employing LC-MS/MS based lipid mediator profiling show that human tenocytes cultured *in-vitro* also produce substantial amounts of the other major prostanoids (PGE_2_ > PGD_2_ > PGF_2α_ > TXB_2_) (52–55). Although PGI_2_ does not appear to be a major product of isolated healthy tenocytes cultured *in-vitro*, stromal cells isolated from diseased human tendons do produce large amounts of PGI_2_ (74). While tendon cells can themselves synthesize prostaglandins, even in the absence of inflammation, mechanically stimulated tenocytes release far greater amounts of PGE_2_ when allowed to interact with MΦ than when cultured in isolation (35). Therefore, cross-talk between infiltrating myeloid cell populations such as PMNs and MΦ with resident tendon cells likely drives the robust prostaglandin response to tendon overuse.

While all major prostanoids were responsive to tendon overuse in the current study, notably they exhibited distinct class-specific temporal responses. Peak PMN infiltration at day 3 of tendon overuse was accompanied by a rapid initial increase in production of PGE_2_ and TXB_2_, while PGD_2_ was simultaneously reduced. Subsequently, PGD_2_ exhibited a more delayed increase from this initial decline to reach concentrations 4-fold above ambulatory controls by day 7 of tendon overuse, at which time the series-J cyclopentenone prostaglandins including PGJ_2_, Δ^12^-PGJ_2_, 15d-PGJ_2_ [which are non-enzymatically derived from PGD_2_ (75)] were also increased. In contrast to PGE_2_, which primarily stimulates inflammation (76), PGD_2_ rather exerts anti-inflammatory actions directly via the DP1 prostanoid receptor, as well as secondary to the formation of downstream cyclopentenone prostaglandins such as 15d-PGJ_2_ which are purported endogenous peroxisome proliferator-activated receptors (PPAR) ligands (77). Thus, overall our data are consistent with prior studies demonstrating a transition from a pro-inflammatory, PGE_2_-dominated eicosanoid profile during the development of inflammation to a more anti-inflammatory, PGD_2_/cyclopentenone-dominated eicosanoid profile during the resolution phase (78).

Earlier studies suggested that in addition to prostaglandins, tenocytes may also release the pro-inflammatory 5-LOX pathway product LTB4 (73, 79). LTB4 biosynthesis involves the initial formation of 5-HpETE via the 5-LOX pathway, which is then converted to leukotriene A4 (LTA4) by the further action of 5-LOX. In cells that express LTA4 hydrolase, LTA4 undergoes subsequent conversion to LTB4. Alternatively, 5-HpETE can undergo further metabolism via reduction or dehydration to 5-HETE and 5-oxoETE, respectively. Both 5-HETE and 5-oxoETE were markedly increased in tendon in response to synergist ablation in the current study, but LTB4 and its downstream inactivation product 12-oxo LTB4 were both below the limits of detection. These data show that heightened mechanical loading of tendon *in-vivo* clearly does increase local biosynthesis of 5-LOX metabolites but question whether LTB4 is a major metabolite produced in tendon. Consistently, recent studies by others utilizing LC-MS/MS also failed to detect LTB4 in isolated human tenocytes *in-vitro* (52–54).

In addition to producing pro-inflammatory leukotrienes, the LOX pathways play important roles in the formation of recently identified SPM families of lipid mediators with pro-resolving actions (80). For example, lipoxin biosynthesis involves the initial production of 15-HETE, a 15-LOX metabolite of n-6 AA, which is then released and taken up by 5-LOX expressing cells (e.g. PMNs) to be converted to LXA_4_ and LXB_4_ (81, 82). In analogous n-3 PUFA generated pathways, 17-HDoHE (a 15-LOX metabolite of n-3 DHA) and 18-HEPE (a CYP metabolite of n-3 EPA) are converted to the D-series resolvins (83) and E-resolvins (62) respectively via the sequential action of 5-LOX. Finally, 14-HDoHE (a 12-LOX metabolite of DHA) serves as the primary intermediate produced during biosynthesis of the most recently identified maresin family of SPMs (42). We show here that tendon contains detectable levels of all four of these major monohydroxylated SPM pathway markers. Tendon overuse markedly increased local concentrations of 15-HETE, 18-HEPE, and 17-HDoHE. On the other hand, 14-HDoHE was unchanged in response to tendon overuse, as were the related 12-LOX metabolites of AA (12-HETE) and EPA (12-HEPE). Overall these data show that, 15-LOX derived SPM biosynthetic pathways are induced in the mechanically overloaded plantaris tendon, like they are in functionally associated muscle (48). In contrast, metabolites of the 12-LOX pathway which are major metabolites produced within functionally overloaded skeletal muscle (48), do not appear to be responsive to heightened mechanical loading of tendon.

We also detected RvD1 in tendon, but its concentration was apparently not influenced by tendon overuse. In contrast, RvD2, protectin D1, and RvD6 were below the limits of detection in tendons from ambulatory rats, but did increase in concentration to become detectable following synergist ablation. Other SPMs including the lipoxins (LXA_4_ and LXB_4_), E-series resolvins (RvE1 and RvE3), and maresins (MaR1) were generally below the limits of detection of our LC-MS/MS assay irrespective of time-point. Overall our data are consistent with recent work showing that isolated human tendon stromal cells cultured *in-vitro* may produce LOX-derived SPMs, in addition to classical pro-inflammatory eicosanoids (52–54). However, similar to the case with muscle tissue (48), tendon homogenates *in-vivo* clearly contain far greater concentrations of primary LOX derived monohydroxylated intermediates in SPM pathways than the bioactive SPMs themselves. This may be attributable to the highly transient nature of mature SPMs, their relative enrichment in the extracellular vs intracellular environment, and/or their naturally low concentrations relative to the limits of detection of the LC-MS/MS assay used here.

We found that plantaris tendons from ambulatory rats were apparently devoid of resident myeloid cells. This finding is consistent with many prior studies that have reported that healthy tendons do not appear to contain a notable resident MΦ population (9, 10, 14–17, 51, 84). In contrast, a recent study described the presence of ‘tenophages’ residing within the dense core of Achilles tendons of healthy ambulatory mice that expressed MΦ lineage markers together with the tenocyte lineage marker scleraxis (85). As a positive control (86), we could easily observe many resident ED2 MΦ (CD68^-^CD163^+^ cells) scattered throughout the perimysium and endomysium of plantaris muscles from these same rats (48). Plantaris tendons did clearly contain resident cells that expressed detectable amounts of mRNA encoding these and other immune cell markers as determined by RT-qPCR. Interpretation of this finding is complicated, however, by prior studies showing that various non-myeloid cell types, including fibroblasts, may also express low levels of common myeloid lineage markers such as CD68 (87). Thus, it currently remains unclear whether a bonafide resident MΦ population exists within the dense core of the tendon proper or rather whether populations of resident tenocytes also express relatively lower amounts of markers commonly used to identify myeloid cells. If such cells do reside in the healthy tendon our data show that either the proteins are not synthesized or are very rapidly degraded resulting in expression levels of these markers far lower than those MΦ that are well-established to reside within skeletal muscle.

Unlike in habitually loaded tendon, we observed a robust peritendinous infiltration of both PMNs and MΦ following synergist ablation surgery. These data are consistent with early studies in which repetitive kicking exercise in rabbits was shown to result in Achilles paratendinitis (13, 88). In a series of later studies inflammatory ED1 monocytes/MΦ (CD68^+^ cells) were also shown to infiltrate peritendinous tissues of the upper limb in response to repetitive reaching/grasping activity in rats, (14–17). While PMNs were specifically localized to the tendon periphery, interestingly we did observe a more delayed increase in MΦ within the dense tendon core in the current study. This finding is consistent with a recent report in which CD68^+^ cells were shown to infiltrate within the core of the Achilles tendon proper following 3-weeks of daily intensive treadmill running in mice (10). To our knowledge, only a single prior study has quantified intratendinous ED2 MΦ (CD163^+^ cells) in response to tendon overuse (14). In this study, repetitive upper extremity reaching and grasping in rats resulted in robust infiltration of palmer and forearm tendons by ED1 MΦ (CD68^+^ cells) between 3-6 weeks of overuse, with no change in ED2 MΦ (CD163^+^ cell) number at this time-point (14). Nevertheless, there was a more modest increase in ED2 MΦ (CD163^+^ cells) in the forelimb tendons by weeks 6-8 of continued overuse (14). Overall, our results following synergist ablation appear most similar to prior studies of intratendinous injection of collagenase, in which only a modest and transient increase in tendon CD163^+^ cells occurred at day 3 post-injury (7, 89).

A single bout of resistance exercise in humans results in a transient increase in serum concentrations of many lipid mediators (57). Many of these same lipid mediators also transiently increase within the exercised musculature of human subjects following an acute bout of muscle damaging (eccentric) contractions, suggesting that injured muscle cells may contribute to the systemic lipid mediator response to exercise stress (58). Consistent with this hypothesis, rodent experimental models of muscle injury were recently found to markedly increase intramuscular lipid mediators (47, 48). However, aerobic exercise, which is generally not thought to inflict substantial muscle damage, was also recently shown to result in marked increases in plasma lipid mediator concentrations (90, 91). The remarkable capacity of mechanically overloaded tendon to produce bioactive lipid mediators identified in the current study suggests that tendon may also be an important and underappreciated cellular source of systemic lipid mediator to exercise (92). Indeed, biopsy samples obtained from human patellar tendons were previously found to express far greater amounts of COX-1 and −2 mRNA when compared to those obtained from the quadriceps muscle (93). Earlier rodent studies also showed that tendons and associated intramuscular connective tissues expressed prostaglandin biosynthetic enzymes much more robustly than the myofibers that make up the bulk of muscle (94). Consistent with these prior studies, we found that the plantaris tendon homogenates analyzed here were greatly enriched in prostaglandins when compared to muscle tissue (48). Overall, these data suggest that muscle-associated connective tissues are likely a key cellular source of bioactive lipid mediators that may serve as autocrine/paracrine signaling molecules between tendon and/or muscle fibroblasts and functionally associated muscle and tendon cells. Similarly, muscle and/or tendon derived lipid mediators may potentially exert cross-organ endocrine actions following their systemic release from the mechanically overloaded musculotendinous unit.

In conclusion, we show for the first time that tendon contains a diverse array of bioactive lipid mediators derived from the COX, LOX and CYP pathways, local biosynthesis of many of which is markedly increased in response to mechanical overuse. When compared to muscle, tendons are greatly enriched in COX and LOX metabolites, but is relatively lacking in products of the CYP pathway. A rapid increase in local concentrations of TXB_2_ and PGE_2_ in mechanically overloaded tendons accompanies peritendinous infiltration of PMNs. The subsequent a more delayed increase in intratendinous anti-inflammatory/pro-resolving mediators is accompanied by a progressive PMN clearance and transition to a predominance of MΦ. Despite this, the SPM response appears insufficient to counteract development of chronic tendon inflammation as evidenced by incomplete resolution of the inflammatory response even at 28 days of continued tendon overuse.

## Author contributions

J.F.M conceived the study. S.V.B and K.R.M and supervised the work. J.F.M and K.B.S designed the experiments. J.F.M and D.C.S performed the experiments. J.F.M and K.R.M analyzed the data. J.F.M prepared the figures and wrote the manuscript with input from all authors.

## Acknowledgments

This work was supported by the Glenn Foundation for Medical Research Post-Doctoral Fellowship in Aging Research (JFM) the University of Michigan Department of Orthopedic Surgery, and the National Institutes of Health (NIH) under the awards R01 (AG050676) (SVB), PO1 (AG051442) (SVB), P30 (AR069620) (SVB) and S10 (RR027926) (KRM).

---

**Supplemental Table 1A:** Cyclooxygenase (COX) metabolite concentration (pmol/g) in overloaded rat plantaris tendons

| Pathway | Substrate | Analyte | Control | Day 3 overload | Day 7 overload | Day 28 overload |
| --- | --- | --- | --- | --- | --- | --- |
| Cyclooxygenase | 20:3n-6 | PGE1 | ND | ND | 5.97 ± 2.54 | 7.20 ± 3.63 |
|  | 20:3n-6 | PGF1a | ND | ND | ND | ND |
|  | 20:3n-6 | 15-keto PGE1 | ND | ND | ND | ND |
|  | 20:3n-6 | 13,14dhPGE1 | ND | ND | 1.72 ± 1.07 | ND |
|  | 20:3n-6 | D17-PGE1 | ND | ND | ND | ND |
|  | 20:3n-6 | Bicyclo PGE1 | ND | ND | ND | ND |
|  | 20:3n-6 | 6-keto PGE1 | ND | ND | ND | ND |
|  | 20:3n-6 | 2,3-dinor PGE1 | 64.87 ± 9.10 | 69.00 ± 11.35 | 83.49 ± 27.00 | 55.29 ± 24.36 |
|  | 20:3n-6 | 19(R)-hydroxy PGE1 | ND | ND | ND | ND |
|  | 20:3n-6 | 15(R)-PGE1 | ND | ND | 6.24 ± 3.03 | 3.38 ± 0.00 |
|  | 20:4n-6 | TXB2 | 35.99 ± 4.39 | 100.40 ± 20.67 | 154.67 ± 28.48 | 184.42 ± 25.55 |
|  | 20:4n-6 | 12-HHTre | 13.26 ± 0.82 | 27.49 ± 4.57 | 57.07 ± 7.58 | 52.89 ± 5.12 |
|  | 20:4n-6 | PGD2 | 56.40 ± 5.13 | 30.31 ± 4.76 | 208.69 ± 64.00 | 219.80 ± 60.36 |
|  | 20:4n-6 | PGE2 | 75.09 ± 6.70 | 257.97 ± 79.88 | 796.02 ± 245.10 | 443.39 ± 69.52 |
|  | 20:4n-6 | PGF2a | 13.48 ± 1.26 | 17.28 ± 2.54 | 36.48 ± 3.94 | 38.20 ± 3.77 |
|  | 20:4n-6 | 6kPGF1a | 63.77 ± 8.77 | 100.08 ± 21.80 | 275.82 ± 54.00 | 226.97 ± 34.45 |
|  | 20:4n-6 | PGJ2 | 5.00 ± 0.59 | 5.36 ± 0.72 | 22.28 ± 4.43 | 15.07 ± 3.57 |
|  | 20:4n-6 | PGA2 | 9.20 ± 1.85 | 11.90 ± 2.45 | 37.64 ± 6.22 | 24.02 ± 4.88 |
|  | 20:4n-6 | 15-keto PGE2 | 3.55 ± 0.43 | 11.64 ± 2.12 | 19.54 ± 3.56 | 13.83 ± 1.68 |
|  | 20:4n-6 | 15-keto PGF2a | 4.26 ± 0.70 | 16.23 ± 4.42 | 43.51 ± 3.56 | 24.55 ± 1.68 |
|  | 20:4n-6 | 13,14dh-15k-PGE2 | 2.83 ± 4.08 | 16.68 ± 4.72 | 41.22 ± 11.59 | 21.71 ± 3.21 |
|  | 20:4n-6 | 13,14dh-15k-PGD2 | ND | 11.75 ± 3.75 | 30.80 ± 5.74 | 16.98 ± 2.39 |
|  | 20:4n-6 | 13,14dh-15k-PGF2a | ND | ND | 2.15 ± 1.02 | 1.25 ± 0.62 |
|  | 20:4n-6 | 11dh-2,3-dinor TXB2 | 3.38 ± 1.37 | 3.51 ± 1.41 | 4.44 ± 2.75 | 6.57 ± 4.19 |
|  | 20:4n-6 | 11-HETE | 128.80 ± 12.51 | 201.81 ± 33.16 | 324.08 ± 30.75 | 369.95 ± 41.71 |
|  | 20:4n-6 | Bicyclo PGE2 | ND | 1.89 ± 0.46 | 2.55 ± 0.79 | 1.17 ± 0.38 |
|  | 20:4n-6 | D12-PGJ2 | ND | 5.41 ± 1.96 | 15.29 ± 7.01 | 14.59 ± 3.42 |
|  | 20:4n-6 | 15d-D12,14-PGJ2 | ND | ND | 1.31 ± 0.67 | ND |
|  | 20:4n-6 | 8-isoPGF2a & 11bPGF2a | ND | ND | 1.54 ± 0.59 | 1.31 ± 0.48 |
|  | 20:4n-6 | 19(R)-OH PGF2a & 20-OH PGF2a | ND | ND | ND | ND |
|  | 20:4n-6 | 19(R)-OH PGE2 & 20-OH PGE2 | ND | ND | ND | ND |
|  | 20:4n-6 | 2,3-dinor TXB2 | ND | ND | ND | ND |
|  | 20:4n-6 | 11dh-TXB2 | ND | ND | ND | ND |
|  | 20:4n-6 | tetranor PGEM | ND | ND | ND | ND |
|  | 20:4n-6 | 6,15-diketo PGFa | ND | ND | ND | ND |
|  | 20:4n-6 | iPF-VI | ND | ND | ND | ND |
|  | 20:5n-3 | TXB3 | ND | ND | 2.03 ± 0.96 | 1.49 ± 0.50 |
|  | 20:5n-3 | PGD3 | ND | ND | ND | ND |
|  | 20:5n-3 | PGE3 | ND | ND | 5.83 ± 2.84 | ND |
|  | 20:5n-3 | PGF3a | ND | ND | ND | ND |
|  | 20:5n-3 | 11dh TXB3 | ND | ND | ND | ND |
|  | 20:5n-3 | 15d-D12,14-PGJ3 | ND | ND | ND | ND |
|  | 20:5n-3 | 11-HEPE | 3.01 ± 0.78 | 3.72 ± 1.55 | 8.44 ± 1.60 | 7.57 ± 1.23 |
|  | Sum |  | 486.40 ± 24.18 | 900.70 ± 172.54 | 2190.70 ± 365.80 | 1754.76 ± 187.05 |

Values are mean ± SEM of 8-12 tendons from 4-6 rats/group. ND = Below limits of detection of the assay.

**Supplemental Table 1B:**
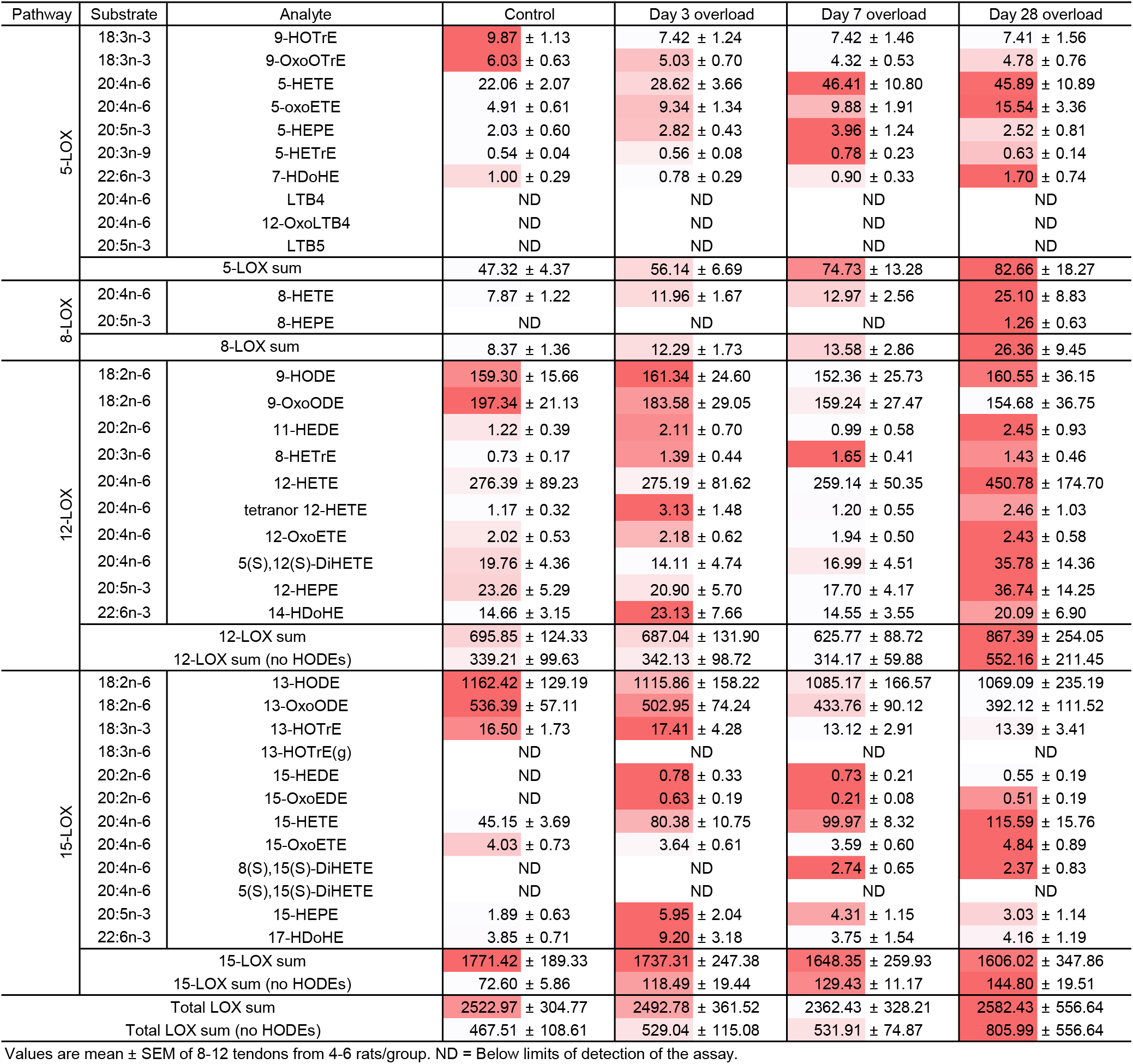
Lipoxygenase (LOX) metabolite concentration (pmol/g) in overloaded rat plantaris tendons

**Supplemental Table 1C:** Epoxygenase (CYP) metabolite concentration (pmol/g) in overloaded rat plantaris tendons

| Pathway | Substrate | Analyte | Control | Day 3 overload | Day 7 overload | Day 28 overload |
| --- | --- | --- | --- | --- | --- | --- |
| Epoxygenase | 18:2n-6 | 9(10)-EpOME | 798.19 ± 83.46 | 741.95 ± 104.39 | 582.41 ± 127.71 | 556.66 ± 100.94 |
|  | 18:2n-6 | 12(13)-EpOME | 687.79 ± 47.87 | 626.89 ± 96.92 | 504.20 ± 128.12 | 403.66 ± 62.71 |
|  | 18:2n-6 | 9,10-DiHOME | 79.79 ± 7.61 | 68.93 ± 10.53 | 63.22 ± 10.77 | 72.02 ± 17.17 |
|  | 18:2n-6 | 12,13-DiHOME | 78.48 ± 6.25 | 69.60 ± 10.00 | 62.11 ± 10.56 | 65.98 ± 11.81 |
|  | 20:4n-6 | 5(6)-EpETrE | 6.63 ± 0.78 | 10.63 ± 1.22 | 10.40 ± 1.60 | 14.85 ± 3.70 |
|  | 20:4n-6 | 8(9)-EpETrE | ND | ND | ND | ND |
|  | 20:4n-6 | 11(12)-EpETrE | 32.29 ± 5.63 | 45.00 ± 5.81 | 40.09 ± 10.00 | 87.48 ± 27.67 |
|  | 20:4n-6 | 14(15)-EpETrE | 16.01 ± 2.44 | 21.46 ± 4.29 | 18.72 ± 5.57 | 43.75 ± 12.39 |
|  | 20:4n-6 | 5,6-DiHETrE | ND | 0.65 ± 0.31 | 0.93 ± 0.39 | 1.56 ± 0.62 |
|  | 20:4n-6 | 8,9-DiHETrE | ND | 0.55 ± 0.18 | ND | 1.04 ± 0.33 |
|  | 20:4n-6 | 11,12-DiHETrE | 2.69 ± 0.89 | 2.48 ± 0.90 | 6.33 ± 0.44 | 8.59 ± 2.35 |
|  | 20:4n-6 | 14,15-DiHETrE | 4.65 ± 0.61 | 6.31 ± 0.79 | 8.20 ± 1.31 | 13.42 ± 2.82 |
|  | 20:4n-6 | 20-HETE | 18.31 ± 3.93 | 22.98 ± 5.38 | 17.45 ± 5.38 | 41.42 ± 12.50 |
|  | 20:5n-3 | 8(9)-EpETE | ND | ND | ND | ND |
|  | 20:5n-3 | 11(12)-EpETE | ND | ND | ND | ND |
|  | 20:5n-3 | 14(15)-EpETE | ND | 3.67 ± 1.17 | ND | 6.53 ± 2.04 |
|  | 20:5n-3 | 17(18)-EpETE | 1.19 ± 0.41 | 1.36 ± 0.48 | 1.27 ± 0.62 | 2.68 ± 0.86 |
|  | 20:5n-3 | 18-HEPE | 1.82 ± 0.99 | 3.52 ± 0.70 | 3.43 ± 0.89 | 4.39 ± 1.25 |
|  | 22:6n-3 | 7(8)-EpDPE | 1.38 ± 0.38 | 1.73 ± 0.37 | 0.76 ± 0.36 | 1.84 ± 0.85 |
|  | 22:6n-3 | 10(11)-EpDPE | 8.06 ± 2.19 | 8.16 ± 1.79 | 5.21 ± 1.34 | 10.60 ± 3.72 |
|  | 22:6n-3 | 13(14)-EpDPE | 4.26 ± 1.13 | 5.45 ± 0.95 | 3.03 ± 0.74 | 5.40 ± 1.72 |
|  | 22:6n-3 | 16(17)-EpDPE | 2.78 ± 0.65 | 4.29 ± 0.63 | 2.30 ± 0.69 | 3.88 ± 1.14 |
|  | 22:6n-3 | 19(20)-EpDPE | ND | 2.45 ± 1.20 | ND | ND |
|  | Sum |  | 1746.31 ± 151.70 | 1648.37 ± 230.92 | 1333.37 ± 275.62 | 1349.65 ± 218.82 |
|  | Sum (no EpOMEs) |  | 102.06 ± 18.55 | 141.00 ± 22.10 | 121.44 ± 24.74 | 251.33 ± 74.69 |

Values are mean ± SEM of 8-12 tendons from 4-6 rats/group. ND = Below limits of detection of the assay.

**Supplemental Table 1D:** Specialized pro-resolving mediator concentration (pmol/g) in overloaded rat plantaris tendons

| Pathway | Substrate | Analyte | Control | Day 3 overload | Day 7 overload | Day 28 overload |
| --- | --- | --- | --- | --- | --- | --- |
| Specialized pro-resolving mediators | 20:4n-6 | LXA4 | ND | ND | ND | ND |
|  | 20:4n-6 | LXB4 | ND | ND | ND | ND |
|  | 20:4n-6 | LXA5 | ND | ND | ND | ND |
|  | 20:4n-6 | 15-epi LXA4 | ND | ND | ND | ND |
|  | 20:4n-6 | 15-oxo LXA4 | ND | ND | ND | ND |
|  | 22:5n-3 | RvE1 | ND | ND | ND | ND |
|  | 22:5n-3 | RvE3 | ND | ND | ND | ND |
|  | 22:6n-3 | RvD5(n-3DPA) | ND | ND | ND | ND |
|  | 22:6n-3 | RvD1 & AT-RvD1 | 32.74 ± 3.98 | 37.77 ± 9.02 | 43.94 ± 11.10 | 38.18 ± 13.79 |
|  | 22:6n-3 | RvD2 | ND | 21.04 ± 10.83 | 45.43 ± 15.68 | 37.73 ± 18.33 |
|  | 22:6n-3 | RvD3 | ND | ND | ND | ND |
|  | 22:6n-3 | AT-RvD3 | ND | ND | ND | ND |
|  | 22:6n-3 | RvD4 | ND | ND | ND | ND |
|  | 22:6n-3 | RvD5 | ND | ND | ND | ND |
|  | 22:6n-3 | RvD6 | ND | ND | ND | 2.92 ± 0.96 |
|  | 22:6n-3 | 8-oxoRvD1 | ND | ND | ND | ND |
|  | 22:6n-3 | 17-oxoRvD1 | ND | ND | ND | ND |
|  | 22:6n-3 | PD1 | ND | ND | 2.58 ± 1.41 | 2.41 ± 1.31 |
|  | 22:6n-3 | AT-PD1 | ND | ND | ND | ND |
|  | 22:6n-3 | 10S,17S-DiHDoHE | ND | ND | ND | ND |
|  | 22:6n-3 | 22-OH-PD1 | ND | ND | ND | ND |
|  | 22:6n-3 | Maresin1 | ND | ND | ND | ND |
|  | 22:6n-3 | 7(S)-Maresin1 | ND | ND | ND | ND |
|  | Sum |  | 34.69 ± 3.98 | 60.67 ± 11.76 | 93.72 ± 24.92 | 82.92 ± 31.06 |

Values are mean ± SEM of 8-12 tendons from 4-6 rats/group. ND = Below limits of detection of the assay.

**Supplemental Table 1E:** Non-enzymatic metabolite concentration (pmol/g) in overloaded rat plantaris tendons

| Pathway | Substrate | Analyte | Control | Day 3 overload | Day 7 overload | Day 28 overload |
| --- | --- | --- | --- | --- | --- | --- |
| Non-enzymatic | 20:4n-6 | 9-HETE | 3.15 ± 0.64 | 3.63 ± 1.30 | 6.07 ± 1.10 | 9.62 ± 3.02 |
|  | 20:5n-3 | 9-HEPE | ND | ND | ND | ND |
|  | 22:6n-3 | 8-HDoHE | 2.42 ± 0.60 | 3.42 ± 0.65 | 2.24 ± 0.63 | 3.64 ± 1.18 |
|  | 22:6n-3 | 10-HDoHE | 3.29 ± 0.66 | 4.93 ± 0.71 | 3.36 ± 0.72 | 5.60 ± 1.82 |
|  | 22:6n-3 | 11-HDoHE | 2.71 ± 0.75 | 3.51 ± 0.85 | 1.92 ± 0.73 | 3.83 ± 1.18 |
|  | 22:6n-3 | 13-HDoHE | 9.09 ± 1.13 | 12.46 ± 1.64 | 11.43 ± 1.42 | 13.82 ± 2.10 |
|  | 22:6n-3 | 16-HDoHE | 7.40 ± 1.24 | 9.17 ± 1.30 | 8.35 ± 1.61 | 11.23 ± 2.75 |
|  | 22:6n-3 | 20-HDoHE | ND | ND | 3.53 ± 1.39 | 4.43 ± 2.17 |
|  | Sum |  | 30.06 ± 4.86 | 40.48 ± 6.73 | 36.99 ± 6.88 | 53.31 ± 14.53 |

Values are mean ± SEM of 8-12 tendons from 4-6 rats/group. ND = Below limits of detection of the assay.

